# High-density surface EMG grid enables non-invasive characterization of intrinsic hand muscles activity

**DOI:** 10.64898/2026.07.10.737782

**Authors:** F. Rizzoglio, V. Darbhe, M. Carvajal, P. Firouzabadi, K.C. Moisio, W. M. Murray, G.L. Cerone, A. Botter, L.E. Miller

## Abstract

Understanding the neuromuscular properties that allow dexterous manipulation of objects remains a major challenge in neurorehabilitation, largely due to the difficulty of characterizing intrinsic hand muscle activity. These muscles are small, densely packed, and anatomically complex, making selective recordings with intramuscular electromyography (EMG) technically demanding and impractical for comprehensive studies. In this work, we present a custom, high-density (HD) surface EMG grid designed to non-invasively capture activity from intrinsic hand muscles from both dorsal and palmar surfaces. We evaluated the quality and spatial selectivity of the recordings by directly comparing them with intramuscular EMG signals obtained from the dorsal and palmar interossei. Surface EMG signals corresponded closely to the intramuscular recordings, with high correlation values for all subjects and tasks. Double differential spatial filtering significantly improved selectivity, although some residual volume conduction remained. The dorsal grid primarily captured dorsal interossei activity, while the palmar grid was more sensitive to lumbrical activation. The palmar interossei recordings were spatially more varied, with the second palmar interosseous predominantly detected on the dorsal grid and the third and fourth on the palmar grid. Together, these results demonstrate that non-invasive HD surface EMG will allow more complete measurement of intrinsic muscle activity, to provide a better understanding of the complex relation between the intrinsic and extrinsic hand muscles during dexterous movements. This basic information will allow refinement of biomechanical hand models and prosthetic devices, and the development of biomimetic brain computer interfaces aimed at restoring natural hand function after neurological injury.

## Introduction

The human hand is the most complex musculoskeletal system in the body, consisting of 27 bones, over 30 muscle-tendon actuators, and extensive soft tissue constraints and linkages, including ligaments and bifurcating tendons [1]. Its remarkable dexterity arises from the action of extrinsic muscles, which originate in the forearm and generate the primary finger and wrist forces, coordinated with the intrinsic muscles, which are located entirely within the hand. The intrinsic muscles, comprising the lumbricals and the dorsal and palmar interossei, play a critical role in controlling the multiple degrees of freedom (DOFs) of the metacarpophalangeal (MCP) and interphalangeal (IP) joints and are essential for fine motor control.

Despite their importance, intrinsic hand muscles remain among the least characterized components of the human musculoskeletal system. Their small size, close spatial arrangement, and deep anatomical location make selective measurement challenging [2]. While activity from many of the extrinsic muscles can be recorded readily using conventional surface electromyography (EMG), recordings from intrinsic muscles typically require intramuscular (IM) electrodes, which provide high selectivity [3], [4]. Even so, targeting these muscles is time-consuming and even painful, requiring multiple needle insertions per finger for full coverage, in an insertion area that is limited by the proximity of the muscles to tendons, nerves, and blood vessels [5], [6]. Consequently, most EMG studies of hand function focus primarily on extrinsic muscles. And yet, intrinsic muscles do have a distinct pattern of neural drive relative to extrinsic muscles [7], [8], evidence comes almost exclusively from static, isometric, single-finger force tasks. Undoubtedly their activity during the dynamic, multi-finger movements typical of natural dexterous manipulation is even less correlated with the extrinsics, but this remains unknown, largely because answering this question would require simultaneous, selective recordings from multiple intrinsic muscles across a broad range of hand configurations and tasks — a scale of data collection that is impractical to achieve with intramuscular electrodes.

A more complete characterization of intrinsic muscle activity could also benefit a wide range of neurorehabilitation and assistive technologies aimed at restoring hand function following neurological injury, including spinal cord injury (SCI), stroke, or upper-limb amputation. In the field of upper-limb prosthetics, recent advances have produced prosthetic hands with numbers of DOFs approaching those of the biological hand [9], [10]. However, intuitive control of these high-DOF prostheses remains a distant goal: despite significant advances in prosthetic hardware, control, including input signals and analytical methods, has not kept pace. Most existing systems still rely on predefined grasp patterns or sequential control strategies [11], [12], [13]. Myoelectric control of upper limb prostheses relies on EMG signals recorded from residual muscles, or even from proximal muscles that have been surgically reinnervated by nerves that would have supplied the distal limb and hand [14], [15], [16], [17], [18]. Biomechanically informed versions of myoelectric control use musculoskeletal hand models and dynamic simulations of hand motion driven by EMG signals, allowing prosthetic control to more closely reflect the mechanics of the natural hand [19], [20]. However, current EMG-driven models overwhelmingly rely on extrinsic muscle inputs alone, largely because comprehensive intrinsic muscle recordings have not been available. Such recordings could be used to refine existing models, by testing whether their inverse dynamics derived estimates of intrinsic muscle activity are consistent with recorded activity. Once validated, these models could run forward dynamics simulations to generate biomechanically informed control signals, either from EMG recorded both intrinsic and extrinsic muscles, or from extrinsic EMG alone, via models trained to approximate the more complete set of muscles in users who lack access to intrinsic signals [21].

A fuller understanding of intrinsic muscle activity would also advance the design of a growing class of intracortical brain computer interfaces (iBCIs) which attempt to provide more biomimetic control. All current iBCI systems for hand control either classify a limited set of postures [22] or decode simple kinematic variables from motor cortical activity [23], [24], [25]. Our group has previously demonstrated that patterns of extrinsic muscle activity can be decoded from motor cortical recordings in individuals with SCI [26]. A muscle-based control system like that described above for a myoelectric interface could extend iBCI control well beyond that of existing kinematic control. In principle, this biometric approach could be extended to a fuller set of intrinsic muscles as well, but doing so requires first knowing what these muscles are actually doing.

To address these needs, we designed a custom high-density (HD) surface EMG grid intended to conform to the anatomy of the intrinsic muscles. This non-invasive approach enables simultaneous recording from multiple muscles on the dorsal and palmar surfaces of the hand without requiring fine-wire intramuscular insertions. We investigated how closely these HD surface EMG recordings correspond to IM EMG signals from select muscles, quantified their selectivity and tested several spatial filtering strategies to increase it.

We show that HD surface EMG signals closely match intramuscular recordings from the dorsal and palmar interossei, albeit with somewhat reduced spatial selectivity that is substantially improved by double differential spatial filtering. The dorsal grid primarily captured activity from the dorsal interossei, the palmar grid predominantly reflected lumbrical activity, and palmar interosseous activity was captured by both grids, in proportions that varied systematically across the digits. Together, these findings demonstrate that HD surface EMG can capture physiologically meaningful, muscle-specific patterns of intrinsic hand activity without the need for invasive intramuscular recordings, providing new opportunities for refined biomechanical models of the hand and supporting the development of prosthetic control systems with myoelectric and brain interfaces aimed at restoring functional hand movements after neurological injury.

## Methods

### Electrode grid design

The HD grid consists of 32 silver electrodes arranged in four strips, each with electrodes laid out on a flexible polyimide substrate with a thickness of 90 µm (Fig. 1A). Each strip is 3.5 mm wide and mechanically independent until they are joined at the proximal end, where the electrical connections to the amplifier are located. Each electrode is ellipsoidal, measuring 2 mm along the major axis and 0.5 mm along the minor axis, with an inter-electrode spacing of 5 mm. The electrode grid is secured to the skin using a double-sided adhesive with cavities aligned with the electrodes. The grid was specifically designed to record from the intrinsic hand muscles: the lumbricals, the dorsal interossei (DI) and palmar interossei (PI). The interossei lie between the metacarpal bones, with the PIs located deeper (from the dorsal surface) than the DIs (Fig. 1A, dark red and green lines, respectively). Although four PIs are described anatomically, that associated with thumb adduction is often rudimentary and was therefore not targeted in the grid layout. Here, 2PI denotes the palmar interosseous that adducts the index finger; 3PI the one for the ring finger; and 4PI for the little finger. The four lumbrical muscles lie closer to the palmar surface and flex the fingers at the metacarpophalangeal joints and extend them at the interphalangeal joints (Fig. 1A, dark blue lines).

**Figure 1.**
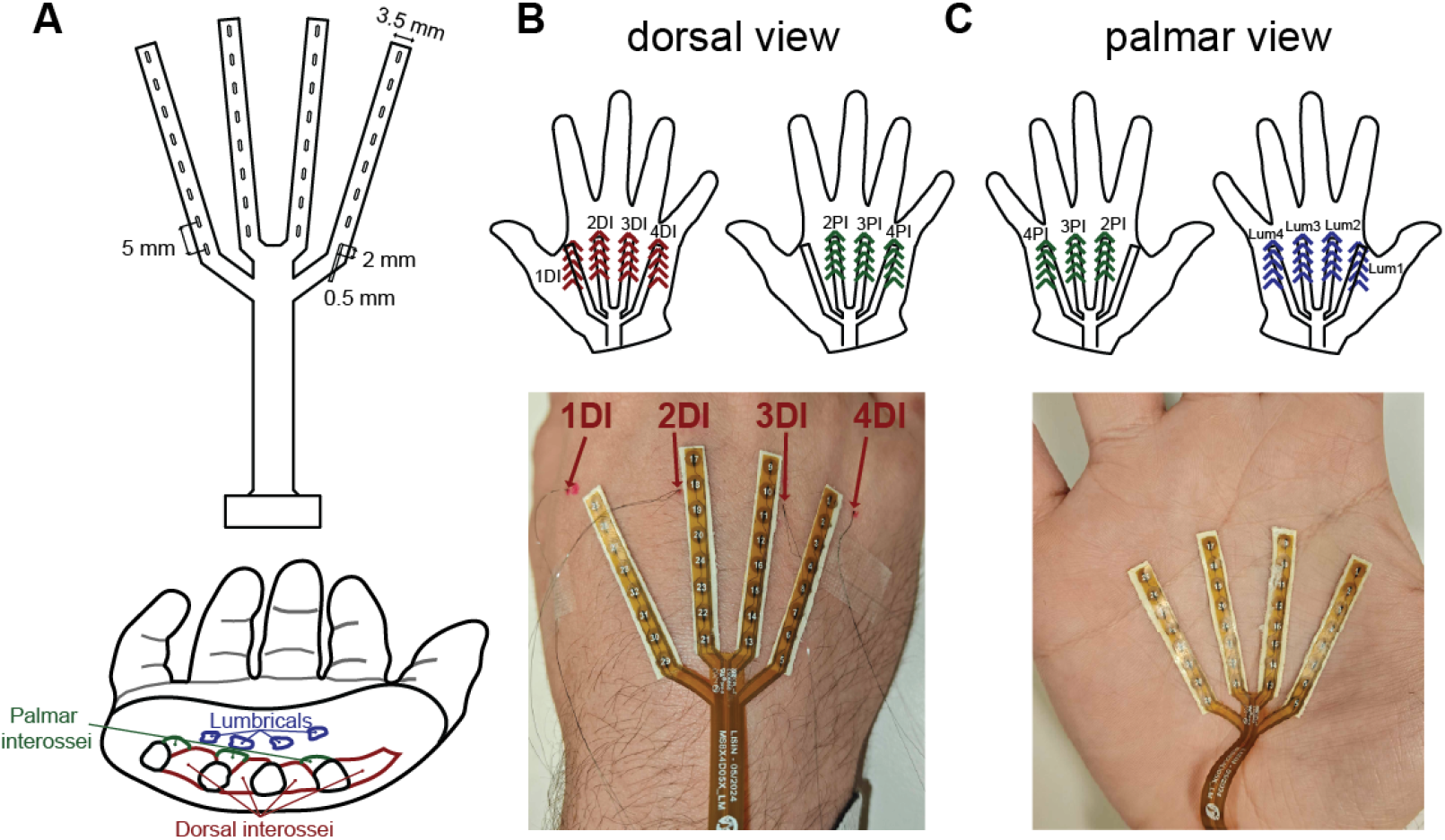
High-density grid for recording intrinsic hand muscles. **A**. Four 3.5-mm wide strips, each containing 8 electrodes and their electrical connections were used to record the activity from the intrinsic hand muscles. The bottom image shows a cross-section of the hand, illustrating the locations of the intrinsic muscles. Most palmar are the lumbricals (dark blue), then palmar interossei (dark green), and then the dorsal interossei (dark red). The metacarpal bones are shown in black. **B**. Placement of the dorsal grid. The top panel shows locations of DIs and PIs using the color scheme from A, along with the grid placement. The bottom panel shows a representative dorsal grid placement and the intramuscular fine wires used to record from the four DIs. Each grid is oriented to align with the underlying muscle, with the IM wires positioned near the distal end. **C**. The palmar grid was aligned with the four lumbricals (dark blue arrows in the schematic). A representative grid placement is shown in the bottom panel.

The conformability of the grid enables its positioning either on the dorsal side of the hand—directly over the four DIs, with the three deeper PIs (Fig. 1B)—or on the palmar side, directly over the lumbricals (Fig. 1C). We expected the dorsal grid to capture activity of both the DIs, anatomically positioned near the dorsal surface, and the PIs, which – despite their name - are closer to the dorsal surface than the palmar surface. In contrast, the lumbricals are the most superficial muscles in the central palm, much closer to the palmar surface, so we expected their activity to be larger in a palmar-paced grid.

### Subjects

We recruited five able-bodied subjects (three males, two females; age 23.6±3.5) for this study. All subjects provided written informed consent according to protocols approved by the Institutional Review Board of the Office for the Protection of Human Participants at Northwestern University.

### Experimental protocol

Our primary objective was to evaluate the spatial distribution of surface HD-EMG activity in the interosseus and lumbrical muscles. Depending on the subject, we recorded surface EMG, intramuscular EMG, or both. For subjects with IM recordings, we inserted fine-wire electrodes into one or more dorsal or palmar interossei. Recording lumbrical activity with IM electrodes is more challenging: fine-wire insertion through the palmar surface is considerably more painful than through the dorsal side, and approaching the lumbricals from the dorsal side is technically much more complex because the insertion needle must pass a considerable depth through additional muscles and tissue. Consequently, we do not have ground-truth IM recordings for the lumbricals.

We recorded both surface and IM EMG in 3 subjects, including one (S1) who visited twice. All subjects performed isometric finger abduction and adduction tasks; subjects with IM recordings (S1-S3) had fine-wire electrodes inserted into targeted interossei, while S4 and S5 — surface EMG only — completed a larger battery of actions including a middle finger tapping task. For subject S1 (first session, denoted as S1a) and subject S2, we inserted fine wires in all four DIs and placed an HD grid on the dorsal side of the hand. For subject S3, we inserted wires in 1DI, 2DI, and 3DI and placed a grid on the dorsal hand. On S1’s second session (noted as S1b), we inserted wires in 1DI, 2DI, and 2PI and placed grids on both the dorsal and palmar sides of the hand. For subjects S4 and S5, we did not make IM recordings. Rather, we conducted a more extensive characterization of the signals recorded from dorsal and palmar HD grids while subjects performed actions designed to differentially activate the DIs, PIs, and the second lumbrical.

All subjects rested their forearm on the table, palm down, with slightly separated fingers, and generated isometric abduction or adduction forces against a force sensor with the specified finger (Fig. 2A). We began by recording maximum voluntary contractions (MVCs) for each finger action. In subsequent trials, subjects started at rest, then slowly ramped the force up to approximately 5–15% of their MVC. After a brief hold, they ramped the force back down to rest. We did not impose strict timing constraints, but we required both the ramp-up and ramp-down phases to last longer than 5 seconds. Subjects S4 and S5 also performed a middle finger tapping action in order to test lumbrical activation. Starting from a resting posture with the MCP flexed such that only the palm and fingertip touched the table, subjects lifted the middle fingertip by extending the proximal (PIP) and distal (DIP) interphalangeal joints (for which the second lumbrical—lum2— is the agonist), with only minimal accompanying MCP extension, then returned the finger to the table in a tapping motion (Fig. 2B). Table I summarizes the recordings obtained across subjects and tasks.

**Table I.**
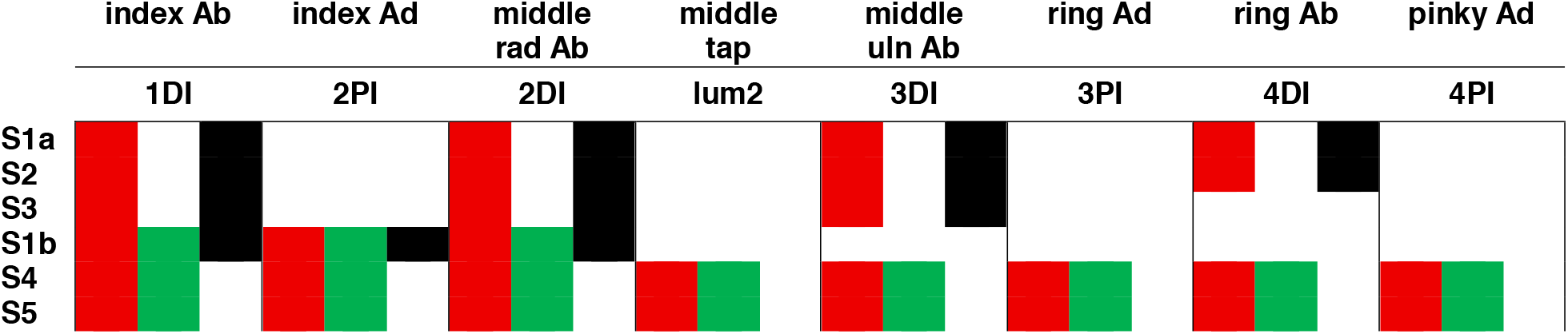
Experimental protocol summary indicating recorded muscles. Subjects listed in rows and task (along with the principal agonist muscle) in columns. Red boxes denote surface HD-EMG recordings from the dorsal side, green boxes indicate HD-EMG recordings from the palmar side, and black boxes represent IM EMG recordings. Note that subject S1 performed two separate sessions, denoted S1a and S1b.

**Figure 2.**
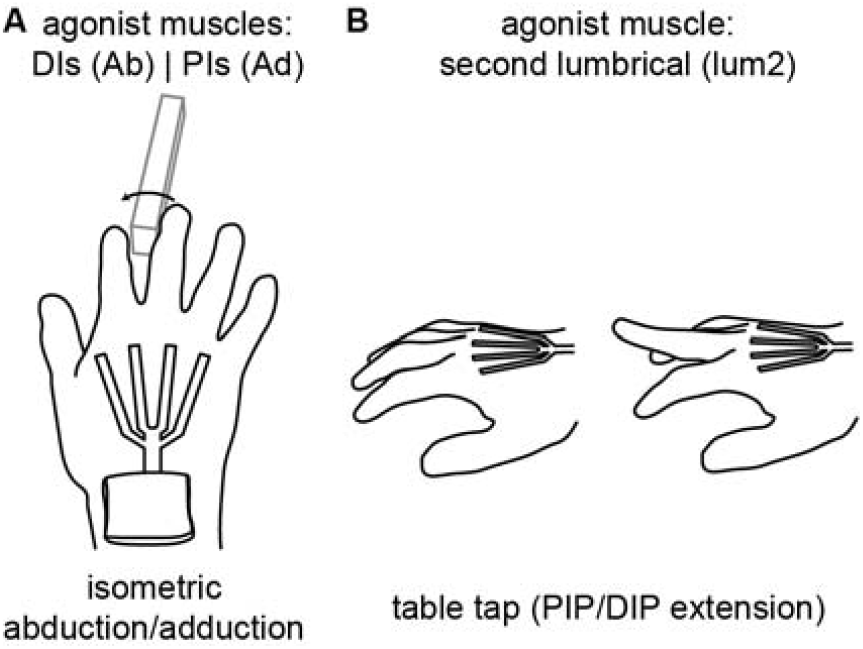
Experimental setup. **A**. In the isometric abduction/adduction task, subjects positioned their hand on a table so they could generate isometric force against a force sensor with a single finger (e.g., radial abduction of the middle finger to activate 2DI). **B**. In the table tapping task, they extended the PIP and DIP joints of the middle finger before returning to the rest position.

At the beginning of each session, we verified correct placement of the IM electrodes by routing the EMG signal to a speaker to listen to EMG modulation while subjects performed an action to selectively activate target muscles (e.g., index finger abduction, in the case of 1DI). We confirmed proper placement when we could clearly hear motor unit activity during activation of the target muscle and no activity during the action expected to silence that muscle (e.g., index finger adduction). If these criteria were not met, we repeated the insertion until we obtained selective activity.

### EMG recording and analysis

We recorded surface EMG signals using a wireless, miniaturized high-density EMG amplifier (MEACS, LISiN, Politecnico di Torino, Italy, and ReC Bioengineering Laboratories, Turin, Italy) [27]. Signals were amplified (192 V/V), band-pass filtered (10 Hz -500 Hz), sampled at 2048 Hz, digitalized with a 16-bit A/D converter, and transmitted via Wi-Fi to a computer for real-time visualization and storage. Although we acquired the signals in a monopolar configuration, we subsequently derived single differential (SD) and double differential (DD) signals during post-processing. We computed the SD signals as the difference between two adjacent electrodes within the same strip, and the DD signals as the difference between two consecutive SD pairs along the strip. Both SD and DD configurations increase spatial selectivity by attenuating common-mode components and reducing volume-conducted activity originating from more distant muscles. In particular, the DD configuration has been shown to further sharpen spatial filtering, thereby limiting cross-talk from adjacent intrinsic muscles [28], [29], [30], [31], potentially improving the localization of motor unit activity.

In addition to the surface EMG recorded with the HD grids, we recorded IM EMG signals from the DIs and the 2PI using Teflon-coated, double-stranded (bifilar) 50 μm wires (California Fine Wire, Grover Beach, CA). The wires were cut without stripping the insulation, exposing only the cross-sectional tip to enhance recording selectivity. Each bifilar pair was inserted into a 30-gauge hypodermic needle, and the tip of the wire was bent to create a small barb to secure the electrode within the muscle after insertion. IM EMG signals were amplified, band-pass filtered (20 Hz–2 kHz), and sampled at 10 kHz using the Bagnoli sEMG system (Delsys, Inc., Boston, MA). The IM electrodes were inserted from the dorsal side of the hand, which is far less sensitive than the palmar skin. Following insertion, the HD grid was positioned such that its strips were placed as close as possible to the inserted fine wires (Fig. 1B).

To facilitate comparison of the two recording modalities, we computed the EMG envelope for both surface and IM recordings by rectifying the raw EMGs and applying a fourth-order Butterworth low-pass filter at 6 Hz. For the HD-EMG signals, this followed derivation of SD and DD signals from the raw monopolar recordings.

We decomposed select surface EMG signals into individual motor unit (MU) spike trains using a validated semi-automated algorithm based on Convolutive Kernel Compensation [32], [33]. Note that we performed MU decomposition only as an exploratory analysis to assess the feasibility of characterizing intrinsic muscle activity at the motor unit level from our recordings, as we did not fully control the experimental conditions required to ensure stable MU recruitment over time (e.g., contraction level was not standardized in percent of MVC, no force feedback was provided to the subject, and the HD-EMG strips were not perfectly aligned with the highly-pinnated muscle fiber directions). We decomposed the monopolar recordings from subjects S1a and S2 during middle and ring finger abduction. We excluded index abduction because, unlike the strips over 2DI, 3DI, and 4DI, the strip over 1DI aligned well only the three most distal electrodes (Fig.□1B). From each identified MU, we computed motor unit action potentials (MUAPs) by spike-triggered averaging an interval from 20□ms before to 20□ms after each discharge, then reconstructed MUAPs for every monopolar, single differential, and double differential configuration. For each action and both subjects, we selected a single MU, with the condition that the Pulse-to-Noise Ratio (PNR) be larger than 27 [34]. For each configuration (monopolar, single differential, double differential), we calculated the RMS value of the MUAP waveform (40□ms window centered on the discharge) at every electrode, yielding a spatial profile of MUAP amplitude along each strip. The per-electrode RMS values were then normalized by the maximum RMS across all electrodes, separately for each MU and configuration. The greater the difference between the normalized RMS on the agonist strip and those on non-agonist strips, the greater the spatial selectivity of that configuration.

## Results

### EMG quality and selectivity

Figure 3 shows example EMG signals from the dorsal side of the hand during a series of different finger abduction forces for S1a. Across all 32 electrodes, monopolar signals (examples in upper row) had an RMS amplitude of 169 ± 65 µV (mean ± SD across electrodes), with a noise floor (EMG amplitude during rest) of 13 ± 6 µV RMS. The corresponding signal-to-noise ratio (SNR) was 17 ± 9. When adjacent signals were differenced (SD configuration), the RMS amplitude decreased to 63 ± 27 µV, with a noise floor of 4 ± 0.9 µV (SNR of 16 ± 7; Fig. 3, second row). When further increasing the spatial filtering through double-differencing, the signal RMS remained unchanged (67 ± 25 µV) and the noise floor slightly increased (6 ± 0.7 µV). Consequently, SNR decreased to 11 ± 4 (Fig. 3, third row).

**Figure 3.**
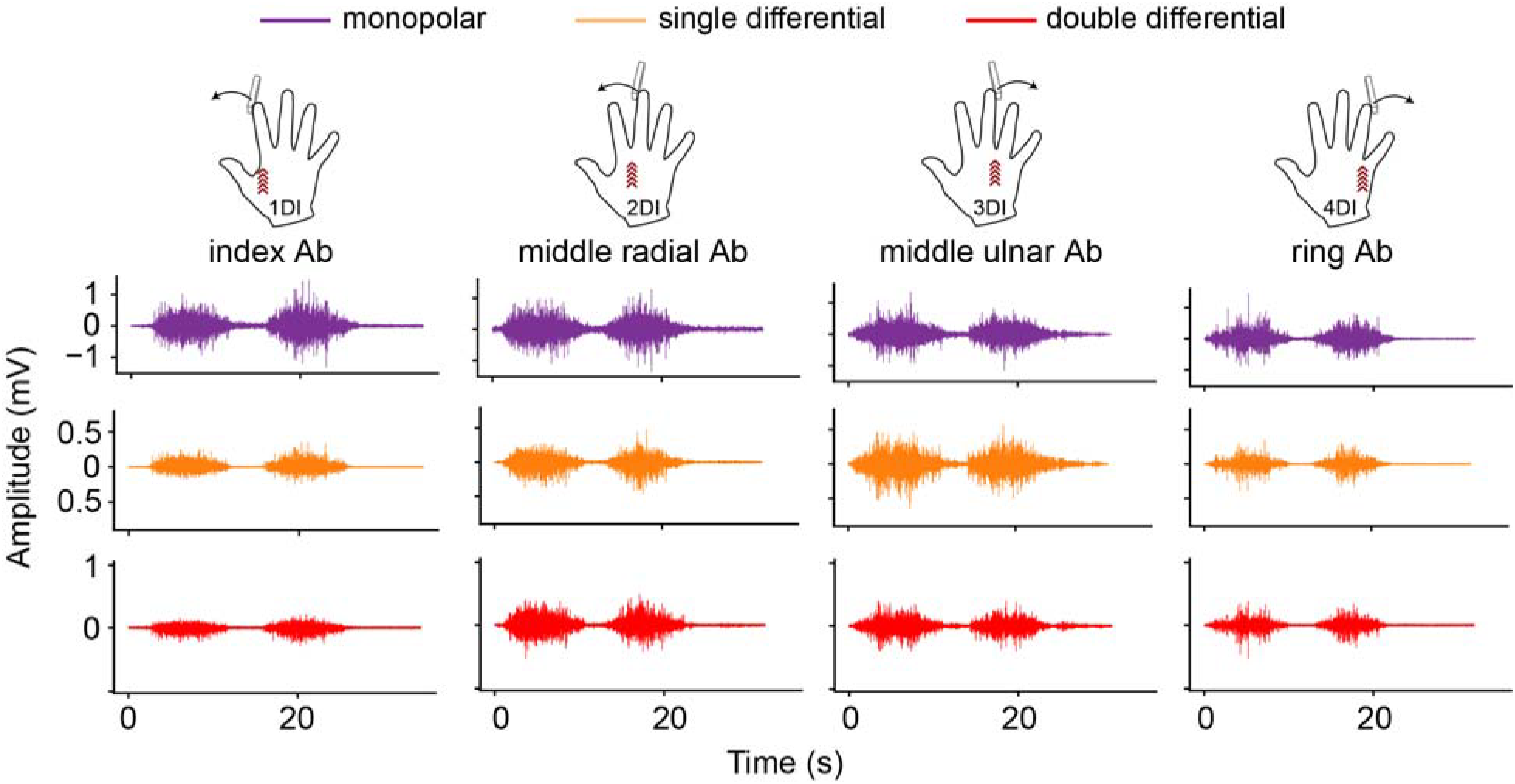
Raw EMG signals with different spatial filtering. For each of the four finger abduction actions, representative single-channel EMG traces for the monopolar (purple, first row), single differential (orange, second row), and double differential (red, third row) configurations are shown from the grid strip positioned over the primary agonist muscle for subject S1a.

We next quantified the spatial selectivity of our recordings. Ideally, each electrode strip should selectively capture activity from the muscle directly beneath it—on the dorsal grid, primarily the dorsal interossei—with minimal contamination from neighboring muscles. To assess detection selectivity, we quantified the similarity between signals recorded from the strip positioned over the agonist muscle and those from adjacent strips by calculating correlation values for each spatial configuration (monopolar, SD, and DD), using the most distal electrode of each strip. We repeated this across the four finger-specific abduction actions. Increasing the spatial filtering tended to decrease the overall amount of correlated activity across the grid (Fig. 4). Going from monopolar to SD reduced the mean correlation from 0.75 ± 0.22 to 0.27 ± 0.21 (Fig. 4, first column); applying DD further decreased it to 0.01 ± 0.18 (Fig. 4, second column). Pairwise correlations for individual subjects are provided in Supplementary Fig.1. In the following sections, we report results using the DD configuration only.

**Figure 4.**
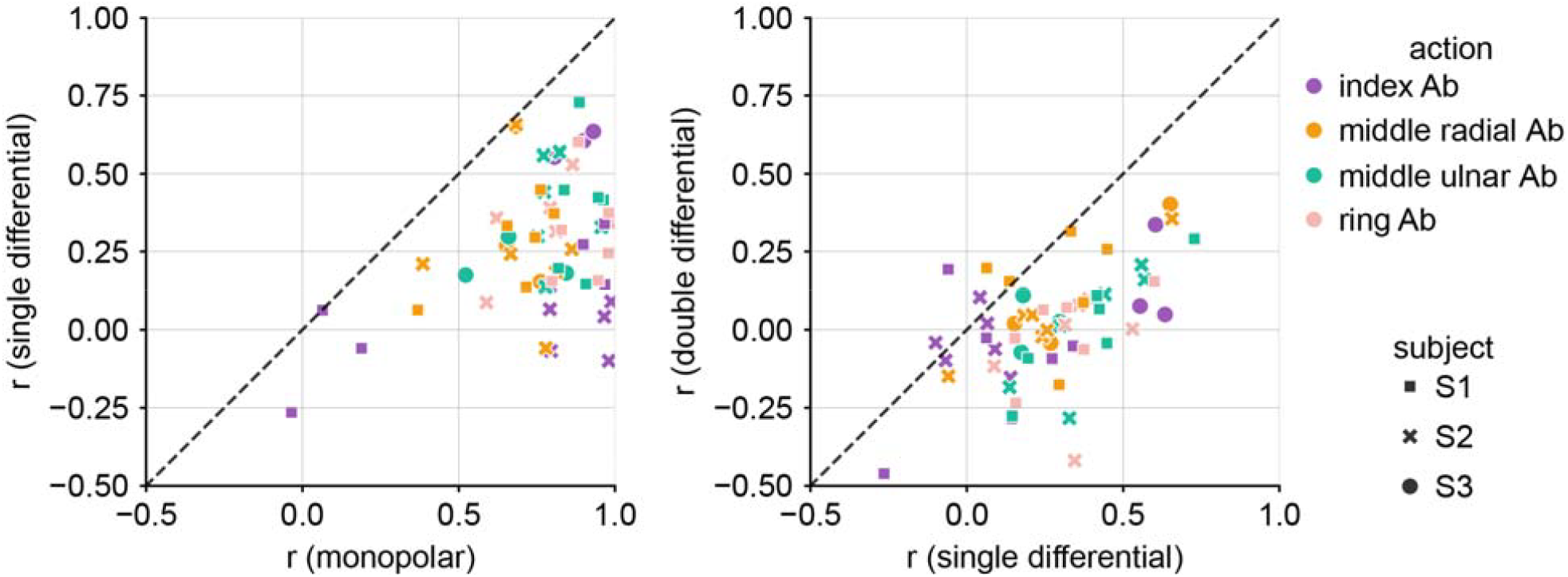
Increasing spatial filtering enhances selectivity. Scatter plots showing Pearson correlation coefficients between the most distal electrode on the strip positioned over the agonist muscle and the most distal electrodes on adjacent strips. Colors denote the particular finger abduction action, while marker shapes represent the three subjects. Each point represents the correlation between the agonist and one of the three adjacent strips for a given action and subject. Higher correlation values indicate lower selectivity. Correlations are shown for monopolar, SD, and DD configurations and directly compared (first column: SD vs. monopolar; second column: DD vs. SD). Greater degrees of spatial filtering progressively reduce correlated activity from adjacent muscles, thereby improving recording selectivity.

### Comparison of surface grid and intramuscular EMGs for dorsal and palmar interossei

After confirming the DD configuration to be the most spatially selective, we next compared those surface recordings with the intramuscular recordings of the underlying muscles. The IM EMG served as ground truth, representing the activity of the muscle in which it was inserted, largely isolated from surrounding muscles. During the task, subjects generated isometric forces to abduct a single finger at the MCP joint against the force sensor. Although the dorsal interosseous associated with each finger was expected to act as the primary agonist for this movement, the extent to which neighboring interossei might also be active was unclear.

The IM recordings confirmed that the dorsal interossei acted as primary agonists during finger abduction (Fig. 5A, black traces in the diagonal panels), and that, for most actions, the envelopes of the surface recordings closely matched those of the IM signals (Fig. 5A, red traces in the diagonal panels), with Pearson correlation coefficients above 0.9 for most pairs. Across all subjects, correlations ranged from 0.58 (1DI, subject S2, Supplementary Fig. 2) to 0.99 (1DI, subject S1a, Fig. 5A), with a mean of 0.91 ± 0.11 (Fig. 5B, diagonal values). The IM recordings also revealed that activation was generally selective for the agonist muscle (Fig. 5A, black traces in non-diagonal black panels), although there was also some activity in non-agonists. For instance, in S1a, 2DI and to a lesser extent 3DI were noticeably active during index finger abduction (for which 1DI is the agonist; Fig. 5A, second and third row, first column).

**Figure 5.**
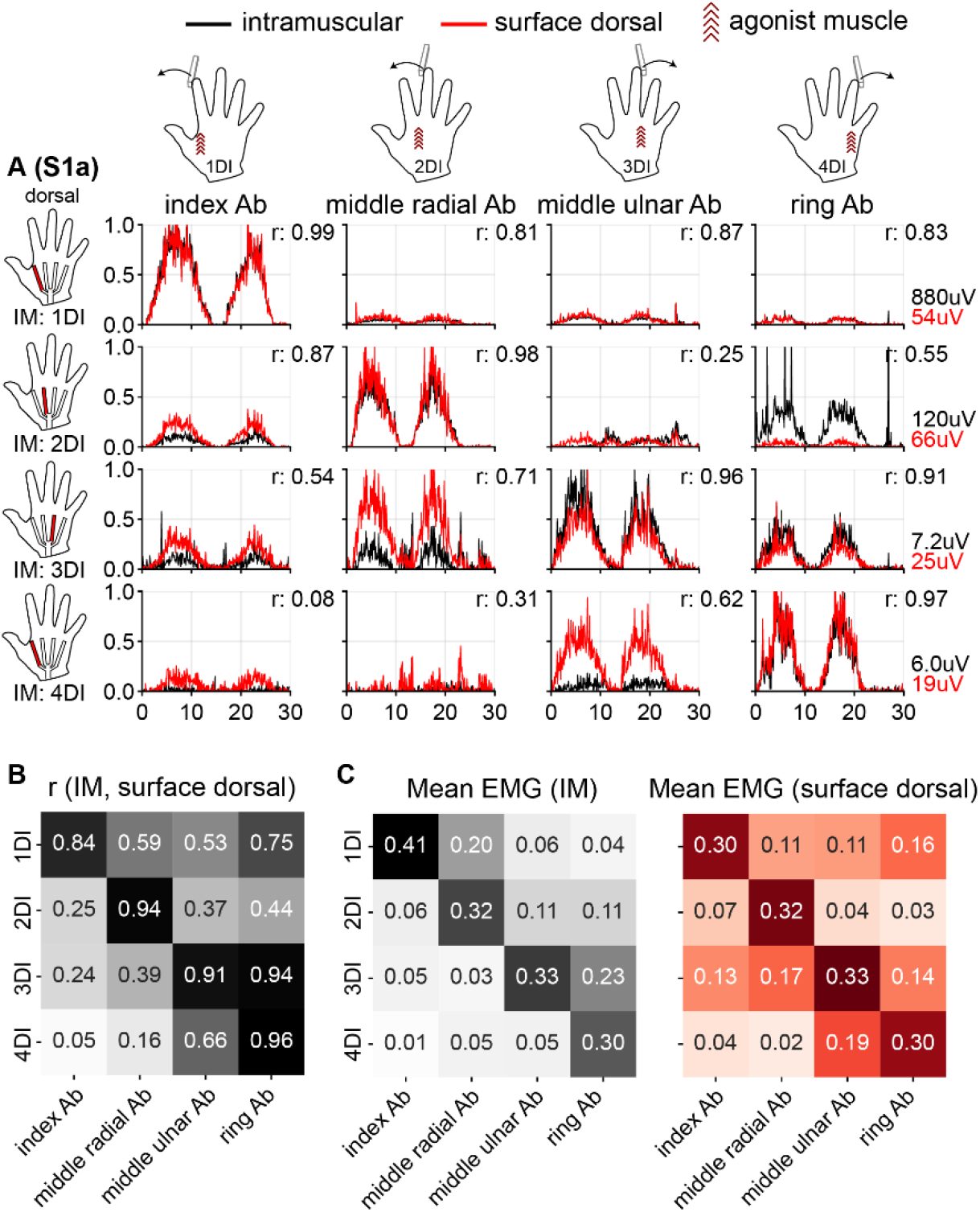
Surface recordings of dorsal interossei have broader spatial activation than IM recordings. **A:** EMG envelopes are shown for each dorsal interosseus muscle during isometric abduction contractions of individual fingers for subject S1a. Black traces represent IM recordings; red traces the double-differential surface signals from the distal end of the dorsal grid. Each column corresponds to a single finger isometric abduction, with recordings from each DI displayed by row. **B:** Pearson correlation between IM and surface EMG envelopes, averaged across subjects S1a, S2, and S3. IM and surface recordings had very similar envelopes in the primary agonist for its respective digit abduction force, with lower values in the non-agonist muscles. **C:** Summary of spatial selectivity across subjects. The left panel presents mean EMG envelope activity for each task from the IM recordings, the right from the dorsal grid. Diagonal values in each matrix indicate the mean activity in the agonist DIs. IM activation was generally selective for the agonist muscle, although low-level activity was sometimes present in non-agonist DIs, likely reflecting stabilization forces. In contrast, the surface EMG envelopes showed a broader spatial activation.

The example IM recordings in Figure 5 were well correlated with the corresponding surface EMG (r = 0.87 and 0.54, respectively), but for both 2DI and 3DI, IM amplitude was smaller than the corresponding surface EMG amplitude. We observed a similar pattern for other conditions, including 3DI during middle finger radial abduction (Fig. 5A, third row, second column), where the correlation with the surface EMG was 0.71. In S2, 1DI was active during both middle finger radial abduction and ring finger abduction, with surface EMG correlations of 0.79 and 0.67, respectively (Supplementary Fig. 2, first row, second and fourth columns). In both S1a and S2, 3DI was active during ring finger abduction (correlations of 0.91 and 0.97 with the surface EMG, respectively), and 4DI was active during middle finger ulnar abduction (correlations of 0.62 and 0.70, respectively). In general, the two types of recordings were well correlated (mean 0.57 ± 0.29, Fig. 5B), but the spatial extent of surface EMG recordings across adjacent strips was consistently broader than the IM recordings, with the non-agonist EMG activity averaging 24% of the agonist one in IM recordings versus 33% for the dorsal surface recordings (Fig. 5C left and right panel, respectively).

The IM recording from 2PI confirmed that it was the primary agonist during index finger adduction (Fig. 6, second row, second column, black trace). In contrast, the 2DI remained largely inactive based on the IM recording (Fig. 6A, third row, second column, black trace). The IM signal from 2PI was minimal during middle finger radial abduction, for which 2DI acts as the agonist (Fig. 6A, second row, third column, black trace). The dorsal grid recorded activity during these actions, with an envelope closely matching the 2PI IM signal during index adduction (r = 0.88; Fig. 6A, second row, second column, red trace), while the palmar grid had no modulation (Fig. 6A, second row, second column, green trace) in any of these actions.

**Figure 6.**
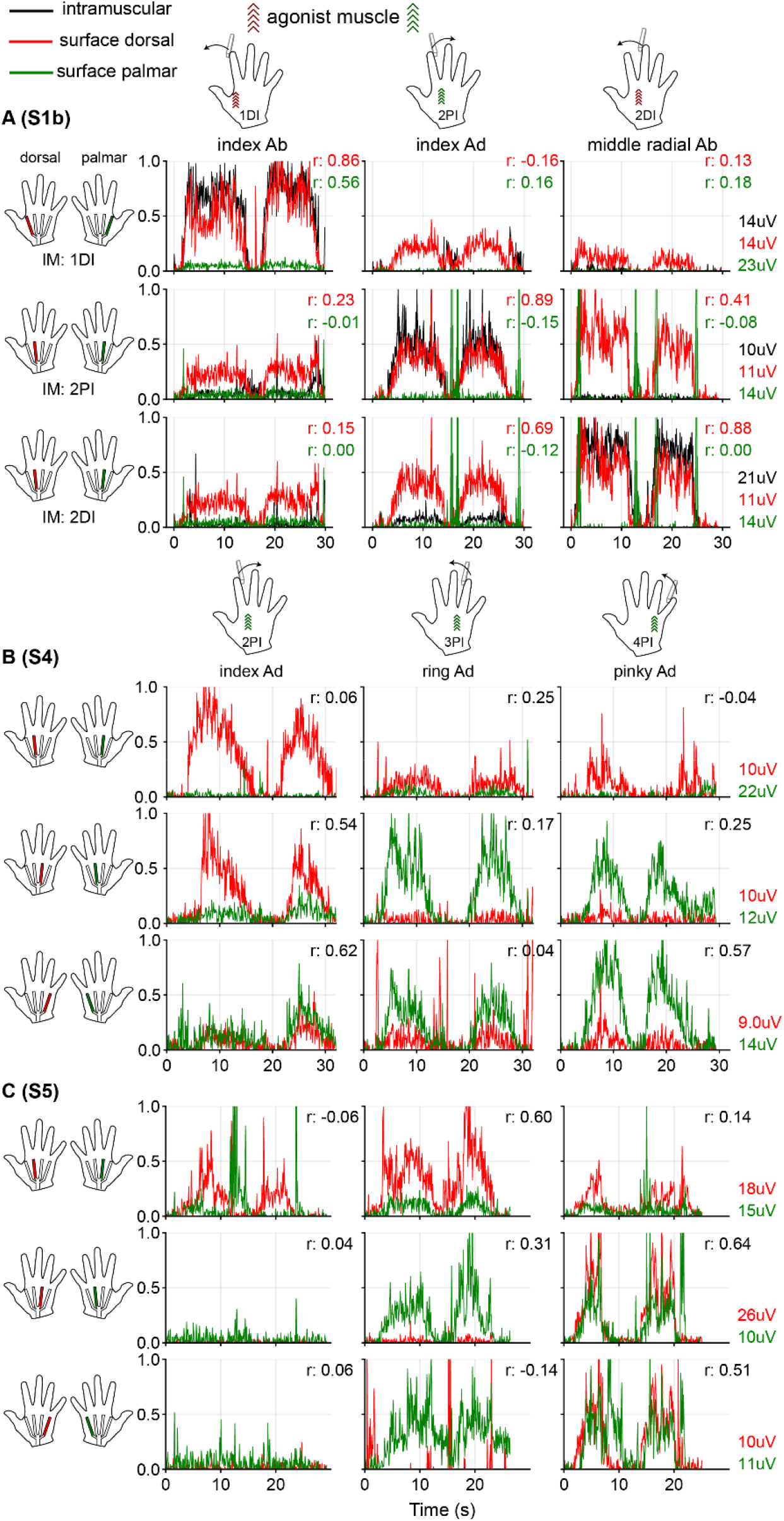
Activity of palmar interossei appeared on both the dorsal and palmar grids. **A:** Activity of the second palmar interosseus is shown together with first and second dorsal interossei during index abduction (left), index adduction (center), and middle radial abduction (right). Black traces represent IM recordings; red and green traces represent double-differential signals from the central channel of the surface grid on the dorsal and palmar aspects of the hand, respectively; the central channel was chosen as it exhibited a slightly higher SNR than the most distal one used elsewhere. Pearson correlations are shown between the IM signal and both dorsal and palmar surface channels. The palmar grid did not record activity from the 2PI during index adduction. **B-C:** Surface recordings during actions in which the palmar interossei act as agonists—index adduction (left), ring finger adduction (center), and little finger adduction (right)—from subjects S4 **(B**) and S5 (**C**). No IM fine-wire electrodes were inserted in these subjects. The palmar grid was more active than the dorsal grid during ring and little finger adduction in both subjects, particularly in S4.

Recordings from S4 and S5, who performed a broader set of adduction tasks without IM recordings, confirmed that, while 2PI activity was detected on the dorsal grid, 3PI and 4PI were detected mostly on the palmar grid. As in S1b, the palmar grid in both subjects was only minimally activated during index adduction, with most of the 2PI activity being detected by the dorsal grid (Fig. 6B–C, first column). However, during ring and little finger adduction—actions in which 3PI and 4PI act as the agonists, respectively—we observed greater activity on the palmar grids compared with the dorsal grid in both subjects (Fig. 6B–C, second and third columns). We also confirmed that the palmar grid did not pick up any dorsal interosseous activity during finger abduction actions where the DIs were the agonists (Supplementary Fig. 3).

Overall, the dorsal surface grid closely captured the activity of the underlying dorsal interossei, with high correlations between surface and the ground truth intramuscular recordings during finger abduction. Although IM recordings were occasionally co-activated with neighboring interossei, the amplitude was smaller than that of the corresponding surface recordings. Activity of the palmar interossei could also be detected by the surface grids, although the relevant grid varied with muscles: 2PI activity was primarily captured by the dorsal grids, whereas 3PI and 4PI activity was more prominent on the palmar grids.

### Lumbrical activity is mostly captured on the palmar grid

We next tested how well the palmar grid captured activity from a representative lumbrical muscle during a task designed to selectively recruit the second lumbrical. When subjects lifted the middle fingertip from the table and fully extended the PIP and DIP joints, we observed substantially greater activity on the palmar grid compared with the dorsal grid (Fig. 7). This pattern was consistent in both S4 and S5 (dorsal RMS during contraction vs. rest: 8.3 vs 5.4 µV for S4; 8.0 vs 5.6 µV for S5; palmar RMS during contraction vs. rest: 18.1 vs 4.8 µV for S4; 26.0 vs 5.4 µV for S5). Overall, these results confirm that the palmar grid is highly sensitive to lumbrical activity.

**Figure 7.**
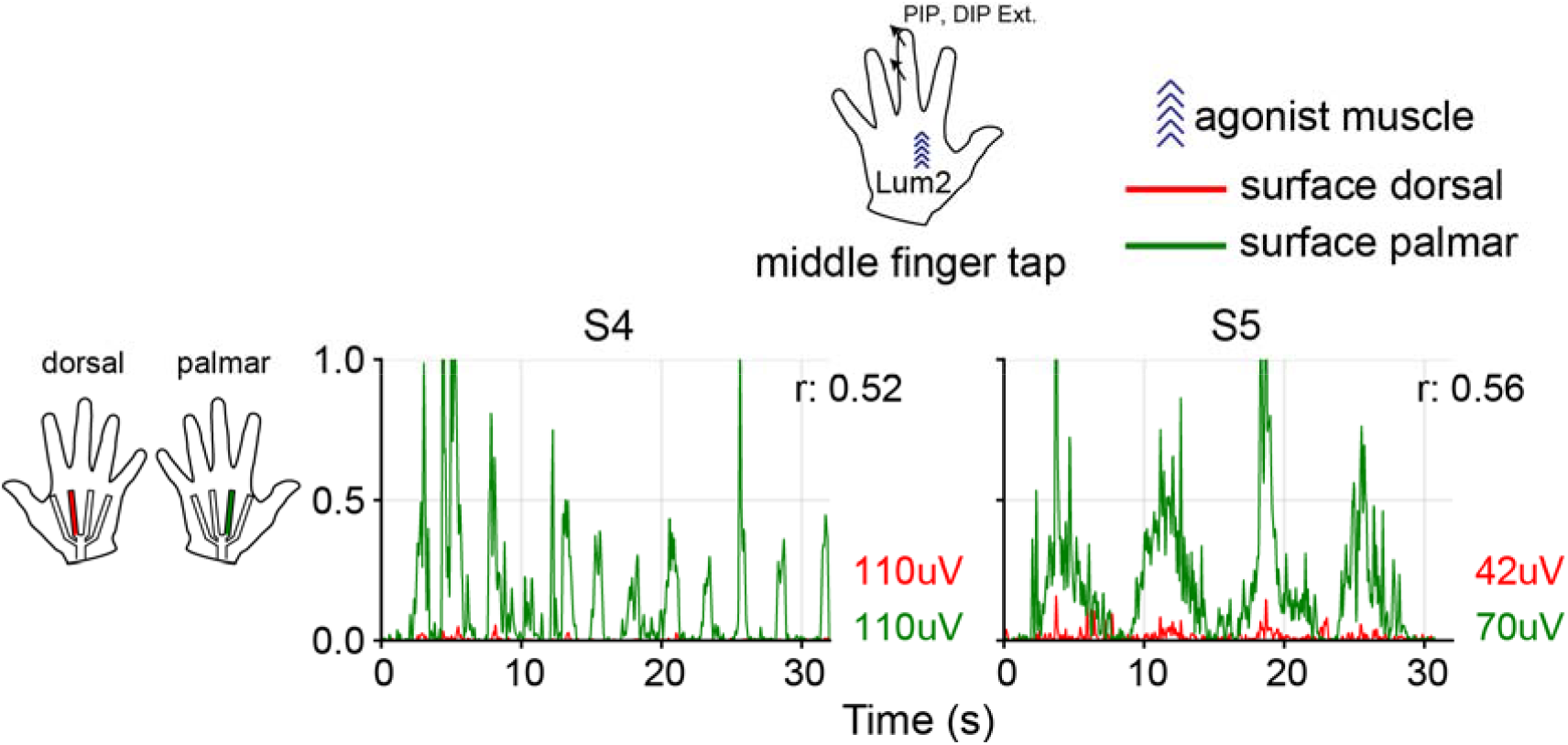
Activity of second lumbrical appeared mostly on the palmar grid. Activity of the second lumbrical recorded by the surface grid on the dorsal (red traces) and palmar (green traces) sides of the hand during the middle finger tapping action for subject S4 (left) and S5 (right). The timing of the taps differed between subjects, with S4 tapping at a faster pace than S5. Pearson’s correlation between the two signals is indicated. Although both grids had modulation related to the movement, the palmar grid captured it with substantially larger amplitude.

### Motor unit decomposition of surface signals

As an exploratory analysis, we tested whether individual motor units could be reliably extracted from surface recordings, and whether spatial filtering conferred a similar improvement in selectivity at the level of individual MUAPs as previously observed at the interference EMG level. Motor unit decomposition confirmed that MUAPs were most prominent over the agonist muscle strip even in monopolar recordings, but that spatial filtering, particularly DD, produced only a modest improvement.

We identified MUs for subjects S1a and S2 during middle and ring finger abduction. Fig.□8A shows an example MUAP from subject S1a during middle finger ulnar abduction for monopolar, single differential, and double differential recordings. In all cases, the MUAP was most prominent on the third strip, which lies over 3DI, the agonist for this action, and differing degrees of spatial filtering progressively limited the MUAP spread to neighboring strips. To quantify this effect, we evaluated the decay of the MUAP amplitude for all tasks and subjects and found that the strip positioned over the agonist muscle consistently had the highest MUAP RMS amplitude, with adjacent strips retaining little amplitude even in monopolar recordings; SD and DD had only modestly lower values in non-agonist strips (Fig.□8B) with the greatest overall decay (i.e., the highest selectivity) for DD. When we repeated the analysis on the raw EMG signals, the substantially higher RMS values for the non-agonist strips in monopolar recordings compared to the corresponding MUAP values, dropped dramatically when moving to SD and DD (Fig. 8C). This is consistent with the strong effect that moving from monopolar to SD had on reducing correlations in raw EMGs across adjacent strips (Fig. 4). The more limited additional RMS reduction from SD to DD is consistent with the increase in noise floor observed with DD (Fig. 3).

**Figure 8.**
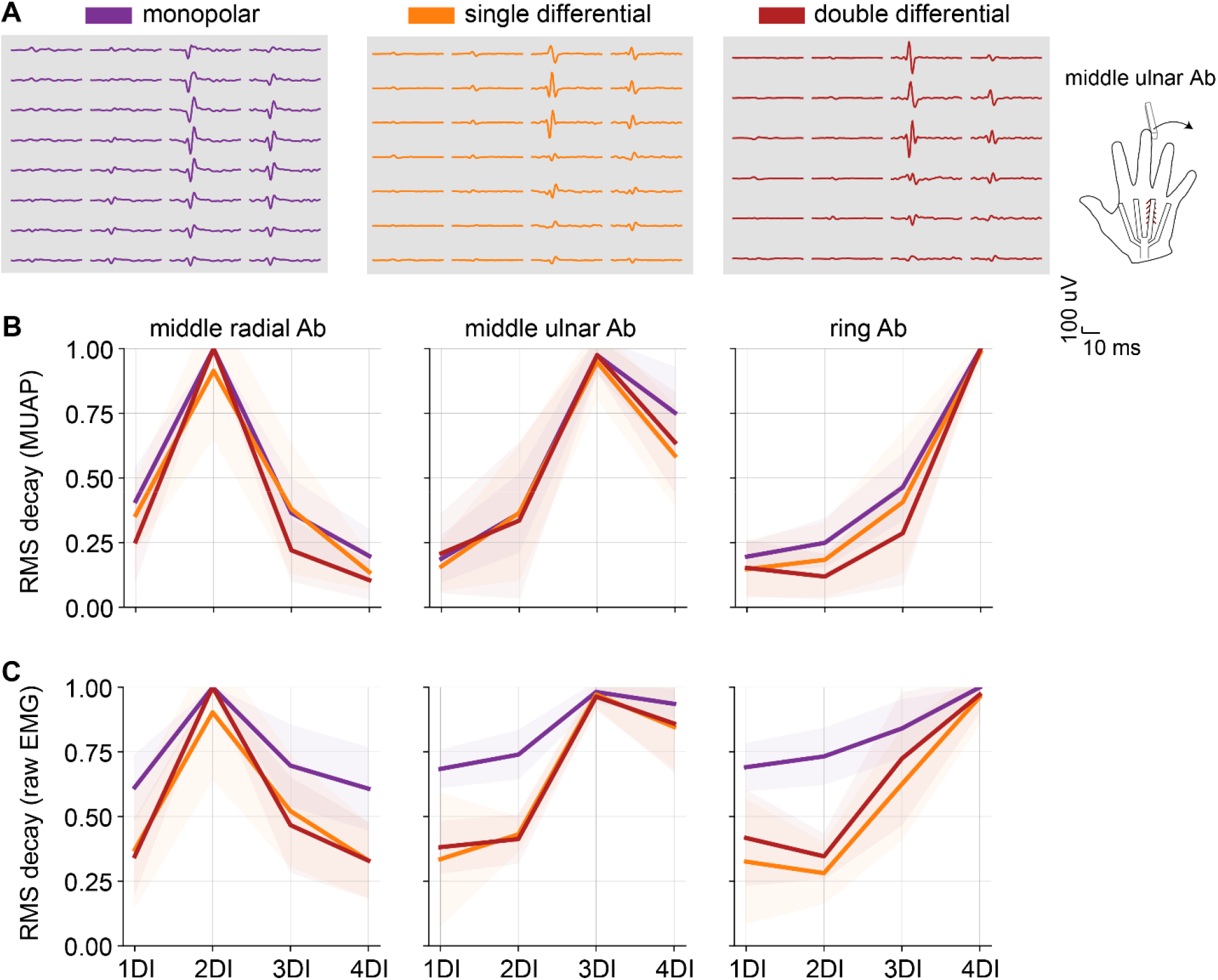
Concentration of motor unit action potentials on the agonist strip was greatest in double differential mode. **A**. Example MUAP from subject S1a during middle finger ulnar abduction (Pulse-to-noise ratio = 29), shown for monopolar (purple, left panel), single differential (orange, center panel), and double differential (dark red, right panel) configurations. The MUAP reached its largest amplitude on the third strip, placed over the agonist (3DI). Its spread to neighboring strips progressively decreased from monopolar to double differential configuration. **B-C**. Percentage of MUAP (**B**) and raw EMG (**C**) total RMS on agonist compared to non-agonist strips for the three spatial configurations. The double differential configuration yielded the greatest concentration of MUAP amplitude on the agonist strip.

## Discussion

This work was motivated by the need to understand intrinsic muscle activity and to integrate it into hand biomechanical models, such as that of Blana et al. [20] for forward-dynamics simulation and model-based control. Recorded EMG signals can serve as proxies for neural excitations, which these models transform into predicted hand movements, and ultimately, into muscle-like control commands for a prosthetic device [20], [35]. Ideally, a fully functionally prosthetic hand, would be driven by information (either measured or inferred) about all relevant extrinsic and intrinsic hand muscles [36]. If, for example, it lacked the intrinsic muscles, the user would be unable to abduct or adduct the fingers or to prevent an intrinsic-minus claw posture [20], [37]. In future work, we plan to use these recorded EMGs in an imitation learning framework similar to those described in [38], [39], where a forward-dynamics, musculoskeletal-driven control policy is trained to reproduce recorded body movements by generating optimal muscle activation patterns. Recorded EMGs can serve not only to validate the physiological soundness of the muscle activity generated by learned control policies, but also to provide additional constraints to reduce the feasible solution space [40], [41].

Our results could also support the development of an intracortical brain computer interface (iBCI) used by individuals with SCI to restore the ability to grasp and manipulate objects. Although considerable progress has been made in using brain signals to control proximal limb movements [42], [43], [44], hand control remains rudimentary. Current iBCIs typically decode kinematic commands from recordings made in the motor cortex, thereby providing only limited, indirect control of grasp force. Some rely on separate control strategies for shaping the hand and regulating contact forces (i.e., a kinematic decoder to pre-shape the hand and a force decoder to regulate contact forces [45], [46]). In contrast, an iBCI based on intended muscle activity [22], [47], [48] could provide a more natural and seamless integration of hand posture, joint torque and stiffness, and contact forces using predicted EMGs as inputs to a musculoskeletal model of the hand. This might better reflect the way the motor cortex normally controls grasping and object manipulation, potentially enabling more dexterous and intuitive hand control. A key requirement for such an EMG-based iBCI would be to pair template muscle activity patterns recorded from able-bodied subjects with motor cortex activity recorded from a participant with tetraplegia who attempts to perform the same actions. This combination would form the basis for observation-based decoder calibration analogous to that used for kinematic iBCIs [42], [49]. This strategy builds on our prior work showing that extrinsic hand muscle activity can be decoded from human motor cortex recordings [26], and extends it to the intrinsic muscles.

These myoelectric and iBCI applications depend on our ability to resolve the activity of individual intrinsic hand muscles with sufficient spatial selectivity. Here, we evaluated how well the activity of these muscles could be characterized with surface recordings, and how spatially selective those recordings were compared to those made with intramuscular wires. The interpretation of the surface recordings is, however, complicated by the mechanical interdependence of the digits caused by soft tissue interconnections, which makes it difficult to determine whether correlated EMG activity across grid strips reflects physiological coactivation or electrical cross-talk. Whenever possible, we therefore used intramuscular recordings as the ground truth of muscle activation.

Based on IM recordings, we noted that although each DI generally acted as the primary agonist during its corresponding abduction action, neighboring interossei often were co-activated to at least a low level during isometric force production (e.g., Fig. 5A, panels at row 2-3, columns 1 and 4), perhaps reflecting a stabilizing role. This physiological co-activation contributed, at least in part, to the spread of correlated activity across adjacent strips of the dorsal grid, particularly evident in monopolar recordings (Fig. 4A). Double differential recordings substantially decreased the correlations (Fig. 4B), though not to the extent of the intramuscular recordings (Fig. 5C), indicating the presence of electrical crosstalk, particularly in the monopolar recordings. Importantly, dorsal interosseous activity remained largely confined to the dorsal grid (Supplementary Fig. 3). The low-level activity during abduction recorded on the palmar grid likely reflects lumbrical activation, recruited to support the DIs during finger abduction. Lumbricals may also have been engaged when subjects extended the PIP and DIP joints to stabilize the finger while generating abduction isometric force, contributing to the recorded palmar activity.

Recordings made from the palmar interossei were spatially more complex. While all the DIs were detected predominantly on the dorsal grid, the distribution of PI activity varied across muscles. 2PI appeared solely on the dorsal grid (Fig. 6A, second column, Fig. 6B-C, first column), whereas 3PI and 4PI were more strongly represented on the palmar grid (Fig. 6B-C, second and third column). 2PI was entirely absent from the palmar grid, likely because adductor pollicis lies above it, but not 3PI and 4PI, as it courses toward the third metacarpal [50]. This anatomical overlap implies that the activity recorded on the two most radial strips of the palmar grid is likely to be strongly confounded by thenar muscle activation, although we did not explicitly test this. While we detected some 4PI activity from the dorsal grid in subject S5 (Fig. 6C, third column), we have no explanation for its absence in S4, or the complete absence of 3PI activity in either subject.

In contrast, the surface recordings from the 2^nd^ lumbrical muscle were more straightforward, with clear representation on the palmar grid (Fig. 7). Taken together, these results suggest that the grids can distinguish most intrinsic muscle activity, with particularly clear selectivity for the dorsal interossei, and for the second lumbrical (and likely the others), which was mainly represented on the palmar grid. We note, however, that the grids have particular limitations for the palmar interossei, with activity patterns that are not confined to a single grid, as the more ulnar PIs may appear on both grids.

Although it was a secondary goal of our study, we were able to extract single motor units from the dorsal interossei, the shape of which remained stable across tasks (Fig. 8). However, unlike Tanzarella et al. [51], we did not observe clear propagation of motor unit action potentials along the strips, perhaps because of the significant pennation angle of the muscle fibers [52]. An additional difficulty was the greater variety of finger muscle contractions in our protocol, which made generating precise force ramps challenging. As a result, MUs were likely recruited and derecruited repeatedly over the course of the contraction, making single motor unit decomposition less accurate.

The design of our high-density EMG grid builds on prior work but incorporates modifications intended to improve coverage of the intrinsic hand muscles [51], [53]. The closest related approach is that of Lara et al. [53], who designed an HD grid of shape similar to ours, from which they could classify a set of hand motions and grasping types. Our grid design incorporates larger spacing between electrode strips and coverage aligned with the muscle bellies, making our grid more suitable for recording EMG activity from individual intrinsic muscles.

