## Supplementary Figures for "High-density surface EMG grid enables non-invasive characterization of intrinsic hand muscles activity"

***Supplementary Information***

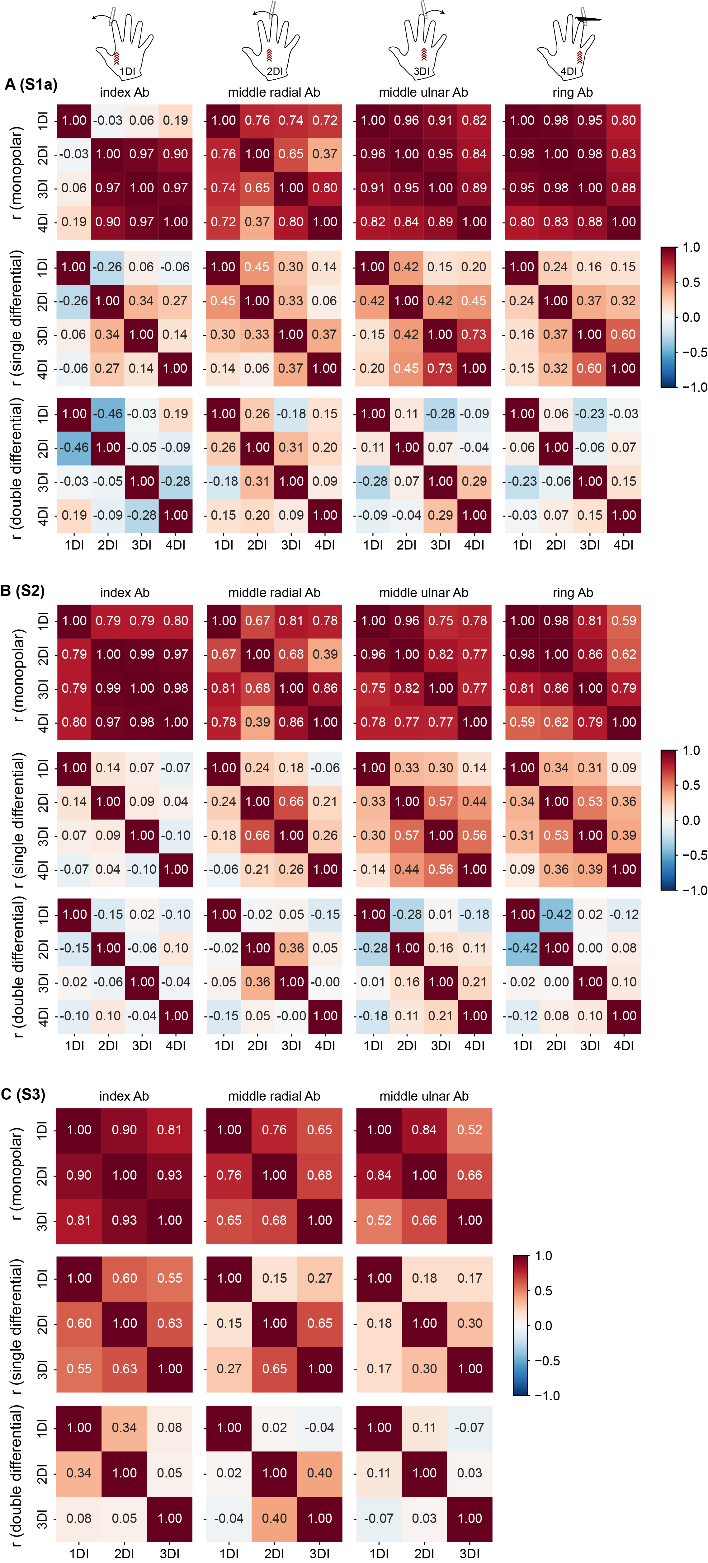

**Supplementary Figure 1.** Pearson correlation matrices of raw surface HD-EMG recordings from the dorsal grid for subjects S1a (**A**), S2 (**B**), and S3 (**C**). In each matrix, rows and corresponding columns represent individual intrinsic muscles. Each panel shows the four isometric finger abductions (columns), while rows show correlation values for the monopolar (first row), single differential (second row), and double differential (third row) configurations. Higher correlation values reflect lower selectivity. Across all subjects, the double differential configuration consistently exhibited the highest selectivity.

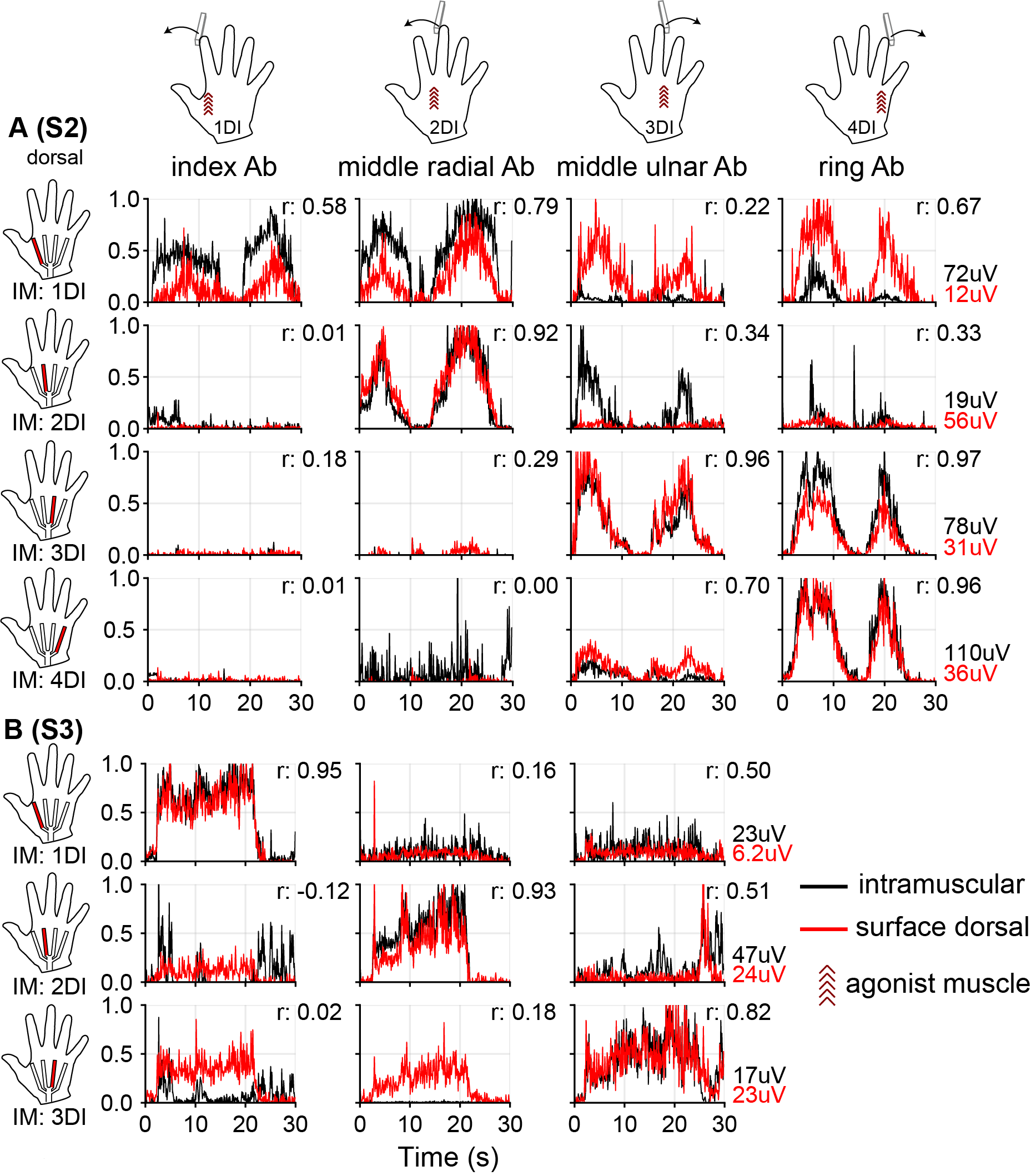

**Supplementary Figure 2.** EMG envelopes for each dorsal interosseus muscle during isometric abduction of individual fingers. Each panel shows data from a different subject: **A** (S2), **B** (S3). Black traces represent IM recordings; red traces the double differential surface signals from the distal end of the dorsal grid. Each column corresponds to a single isometric finger abduction, with recordings from each DI displayed by row.

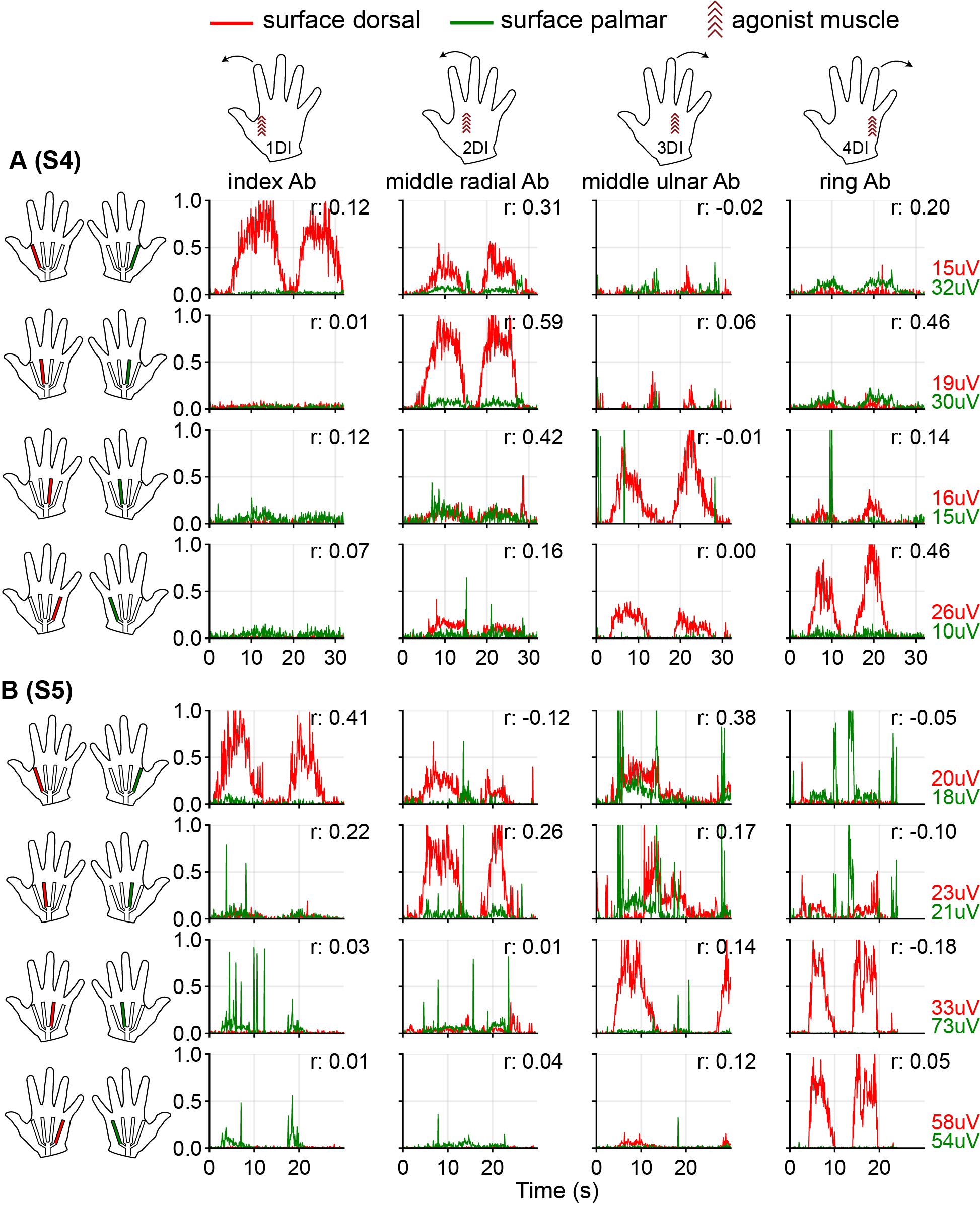
**Supplementary Figure 3.** EMG envelopes during isometric abduction of individual fingers recorded from the dorsal and palmar grids. Each panel shows data from a different subject: **A** (S4), **B** (S5). Red traces show double-differential surface signals from the distal portion of the dorsal grid, and green traces show signals from the distal portion of the palmar grid. Each column represents a single finger abduction task, while rows display recordings from each grid strip (ordered from top to bottom, ranging from most radial to most ulnar for both grids). The dorsal grid consistently captured activity from the agonist dorsal interossei, whereas the palmar grid had little to no detectable activity in S4 and only minimal activity in S5.
